# Familial ALS-associated *SFPQ* variants promote the formation of SFPQ cytoplasmic aggregates that reduce surface AMPA receptor expression in primary neurons

**DOI:** 10.1101/2022.03.10.483757

**Authors:** Jocelyn Widagdo, Saumya Udagedara, Nishita Bhembre, Jing Zhi Anson Tan, Lara Neureiter, Jie Huang, Victor Anggono, Mihwa Lee

## Abstract

SFPQ is a nuclear RNA-binding protein that is involved in a wide range of physiological processes including neuronal development and homeostasis. However, the mislocalization and cytoplasmic aggregation of SFPQ are associated with the pathophysiology of amyotrophic lateral sclerosis (ALS). We have previously reported that zinc mediates SFPQ polymerization and promotes the formation of cytoplasmic aggregates in neurons. Here we characterize two familial ALS (fALS)-associated *SFPQ* variants, which cause amino acid substitutions in the proximity of the SFPQ zinc-coordinating center (N533H and L534I). Both mutants display increased zinc-binding affinities, which can be explained by the presence of a secondary zinc-binding site revealed by the 1.83Å crystal structure of the human SFPQ L534I mutant. Overexpression of these fALS-associated mutants significantly increases the number of SFPQ cytoplasmic aggregates in primary neurons. Although they do not affect the density of dendritic spines, the presence of SFPQ cytoplasmic aggregates causes a marked reduction in the levels of the GluA1, but not the GluA2 subunit of AMPA-type glutamate receptors on the neuronal surface. Taken together, our data demonstrate that fALS-associated mutations enhance the propensity of SFPQ to bind zinc and form aggregates, leading to the dysregulation of AMPA receptor subunit composition, which may contribute to neuronal dysfunction in ALS.

## INTRODUCTION

Splicing factor proline-and glutamine-rich (SFPQ) is an RNA-and DNA-binding protein that is involved in many aspects of RNA biogenesis as well as DNA damage repair [1, 2]. It is ubiquitously expressed in most tissues and cell types and has been implicated in a wide range of physiological functions, including neuronal development [1, 3-5]. SFPQ belongs to the Drosophila behavior human splicing (DBHS) protein family together with two paralogs, non-POU domain-containing octamer-binding protein (NONO) and paraspeckle component 1 (PSPC1). The DBHS family proteins share a high sequence similarity within the central DBHS domain which comprises two RNA-recognition motifs (RRMs), a NonA/paraspeckle (NOPS) domain, and a long coiled-coil domain (**Fig. 1A**). It has previously been shown that SFPQ homo- and hetero-dimerizes via the central DBHS domain and polymerizes via the extended coiled-coil domain, which is critical for its nuclear functions, such as transcriptional regulation and paraspeckle formation [6-8]. Although SFPQ primarily functions within the nucleus, there is increasing evidence for cytoplasmic functions of SFPQ, which include the regulation of neuronal RNA transport [9-11]. These observations indicate that the correct balance in the nucleocytoplasmic distribution of SFPQ is critical for neuronal development and homeostasis. The cytoplasmic aggregation and mislocalization of RNA-binding proteins (RBPs) are emerging hallmarks of neurodegenerative diseases including amyotrophic lateral sclerosis (ALS) [12, 13]. These RBPs are best exemplified by trans-activation response element DNA-binding protein 43 (TDP-43) and fused in sarcoma (FUS), both of which predominantly function in the nucleus under normal conditions but are mislocalized and aggregated in the disease state [13, 14]. Recent evidence has also demonstrated abnormal cytoplasmic accumulation and loss of the nuclear pool of SFPQ in ALS [15, 16]. However, the precise molecular mechanisms that underpin these pathological changes, and their effects on neuronal functions are not well understood.

**Fig. 1.**
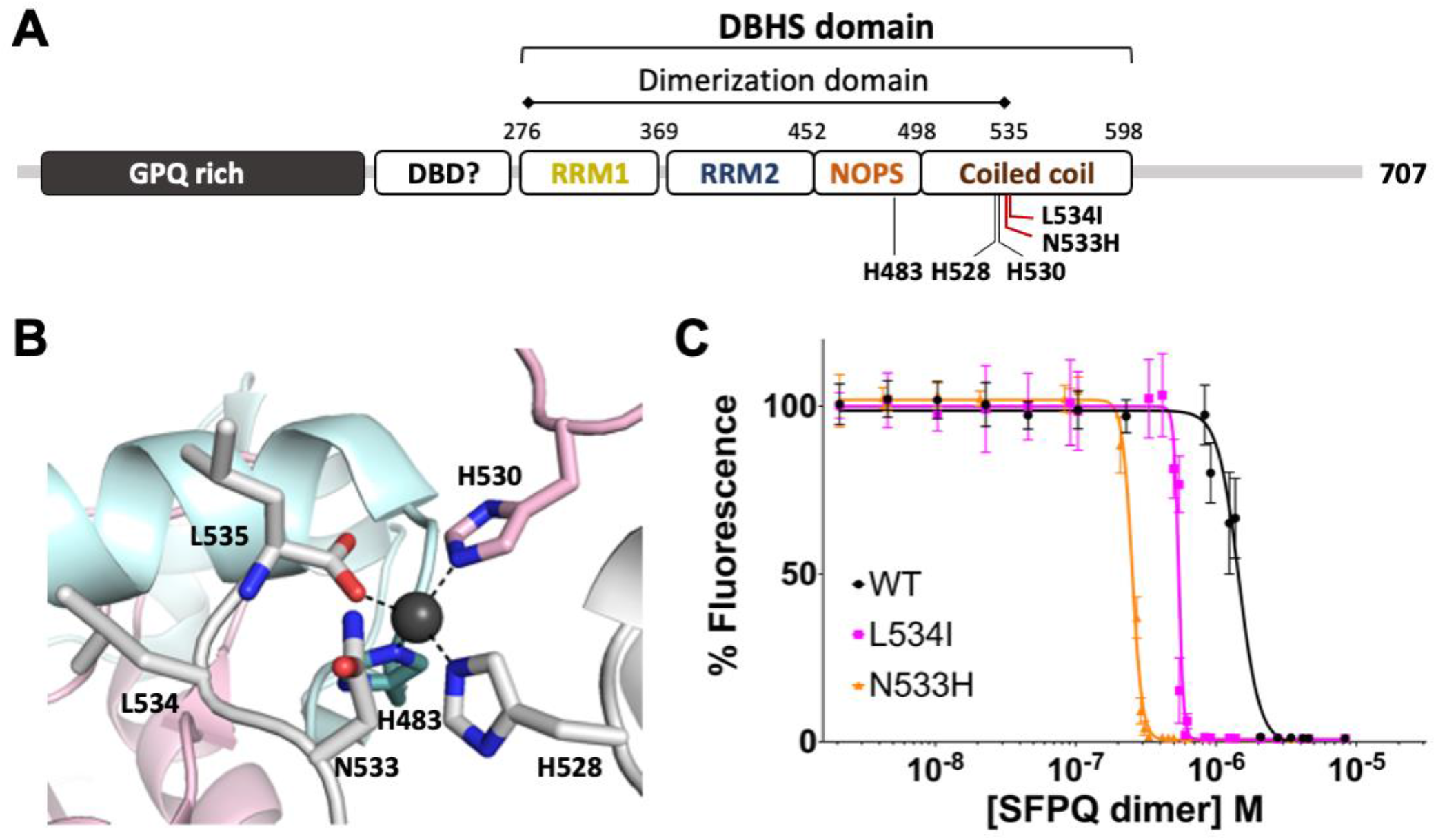
SFPQ N533H and L534I bind to zinc with higher affinities than the WT protein. **A** Schematic domain organization of human SFPQ, depicting the positions of the zinc-coordinating residues (H483, H528, and H530) and the reported fALS-associated mutations, N533H and L534I. **B** Crystal structure of human SFPQ with a close-up view of the zinc-coordinating center (PDB code 6OWJ [19]) with the side chains of fALS-associated mutation residues (N533 and L534) displayed in stick presentation. **C** The N533H and L534I fALS-associated SFPQ mutants display higher zinc-binding affinities measured by the Zn^2+^ indicator, Fluozin-3 compared to the WT protein. Data represent mean ± SD from three independent experiments.

Altered metal homeostasis and increased oxidative stress have consistently been proposed as central features of neurodegeneration [17, 18]. Zinc is a transition metal that is selectively stored in, and released from, the presynaptic vesicles of some neurons, with its dysregulation exerting detrimental effects on these cells [18]. We recently discovered that zinc binds to SFPQ and induces SFPQ polymerization *in vitro*, as well as promoting the formation of SFPQ cytoplasmic aggregates in primary neurons [19]. This study supports the notion that the dysregulation of zinc in the brain triggers an imbalance in the nucleocytoplasmic distribution of SFPQ, with several studies reporting an elevated level of zinc in the serum and cerebrospinal fluid of ALS patients [20-22].

Here, we focused on two recently identified missense mutations of SFPQ in familial ALS (fALS; N533H and L534I), both of which cause morphological abnormalities in the axons of motor neurons in the zebrafish [16]. Interestingly, these mutations are located in close proximity to the intermolecular zinc-coordinating center of SFPQ, which is composed of His-483, His-528, His-530 and the carbonyl oxygen of Leu-535 [19]. We therefore tested the hypothesis that these fALS-associated mutations enhance the zinc-binding affinity of SFPQ, thereby increasing its propensity to aggregate in the cytoplasm of disease-affected neurons. We also investigated the effects of SFPQ N533H and L534I mutants on the structure and function of excitatory synapses by examining the density of dendritic spines and the levels of surface α-amino-3-hydroxy-5-methyl-4-isoxazoleproprionic acid (AMPA)-type glutamate receptors (AMPARs) in primary rat cortical neurons.

## METHODS

### DNA constructs

The construction of pCDF11-SFPQ-276–535, pGEX6p1-SFPQ-276–598, and pGEX6p1-RXRα-DBD (residues 130-228) has been described elsewhere [7, 19]. The N533H and L534I mutations in the pEGFP-SFPQ and pGEX6P1-SFPQ-276–598 constructs were generated with the Q5 site-directed mutagenesis kit (New England Biolabs). All constructs were verified by DNA sequencing. Plasmid DNAs encoding the myc-tagged GluA1 and GluA2 subunits of AMPARs were gifts from Prof. Richard Huganir and have been described previously [23, 24].

### Protein expression and purification

His6-tagged SFPQ-276–535 L534I was expressed and purified by the method previously reported for wild-type (WT) His-tagged SFPQ-276–535 [7]. The procedures for the expression and purification of GST-tagged RXRα-DBD-130–228 (encoding amino acid residues 130 – 228 of human RXRα) were performed according to a previous report [25]. GST-tagged SFPQ constructs (SFPQ-276–598 WT, SFPQ-276–598 N533H, and SFPQ-276–598 L534I) were expressed and purified by the method previously reported for GST-SFPQ-276–598 WT [19]. Following the cleavage of the GST-tag, recombinant proteins were then eluted from a HiLoad 16/600 Superdex 200 pg column (GE Healthcare) equilibrated with 20 mM Tris-HCl (pH 7.5), 300 mM NaCl. Final protein samples were concentrated to 4 – 8 mg/ml, snap-frozen using liquid nitrogen and stored at -80°C.

### Zinc-binding assay

Due to the formation of infinite polymers and protein precipitation upon zinc binding, we were not able to measure the binding affinity of SFPQ to zinc directly [26]. Instead, measurements of the zinc-binding affinities of the SFPQ-276-598 WT and mutant proteins (SFPQ-276–598 N533H and SFPQ–276-598 L534I) were achieved with the Fluozin-3 competitive zinc-binding assay as previously described [19, 27]. Briefly, competition by ethylenediaminetetraacetic acid (EDTA)-treated SFPQ proteins in 20 mM MOPS (pH 7.0), 250 mM NaCl for Zn(II) binding was assessed by monitoring the decrease in the fluorescence of 150 nM Fluozin-3-Zn(II) with an excitation wavelength of 485 nm and emission wavelength of 520 nm in response to increasing SFPQ protein concentrations. The data were analyzed using the equation, log_10_[inhibitor] versus response – variable slope, in Prism (GraphPad Software) to determine the IC_50_ value for zinc-binding.

### Crystallization and X-ray diffraction data collection

Crystals of the Zn-SFPQ-276–535 L534I complex were grown using the same strategy as for SFPQ WT 276-535 in complex with Zn(II). Briefly, SFPQ-276–535 (2.5 mg/ml) and RXRα-DBD-130–228 (1 mg/ml), both in 20 mM Tris-HCl (pH 7.5), 500 mM NaCl were mixed at a molar ratio of 1:1 and incubated for one hour before being concentrated to 15 mg/ml. The crystals were grown using the hanging-drop vapor diffusion method at 20°C by mixing 2 μl of SFPQ-RXRα-DBD (7.5 mg/ml) with 2 μl of reservoir solution [0.1 M MES (pH 6.0), 0.2 M calcium chloride, and 12% (w/v) PEG 4000] and equilibrating against 0.5 ml reservoir solution. Prior to cryo-cooling, crystals were successively transferred to artificial reservoir solutions containing 20% ethylene glycol in 10% increments. Diffraction data were recorded on beamline MX2 at the Australian Synchrotron [28] at a wavelength of 0.954 Å at 100 K. Additional datasets were collected for metal identification at low- and high-energy remote wavelengths of 9560 eV and 9760 eV, respectively, at 100 K. The data were processed with XDS [29], and merged and scaled with AIMLESS [30]. Crystals belong to space group *P*2_1_ with unit cell parameters of *a* = 61.6, *b* = 62.7, *c* = 67.8 Å, and *β* = 96.1°, similar to those of the WT crystals [19]. Data collection and merging statistics for the native data set are summarized in **Table 1** and those for the data sets collected for metal identification in **Supplementary Table S1**.

**Table 1.**
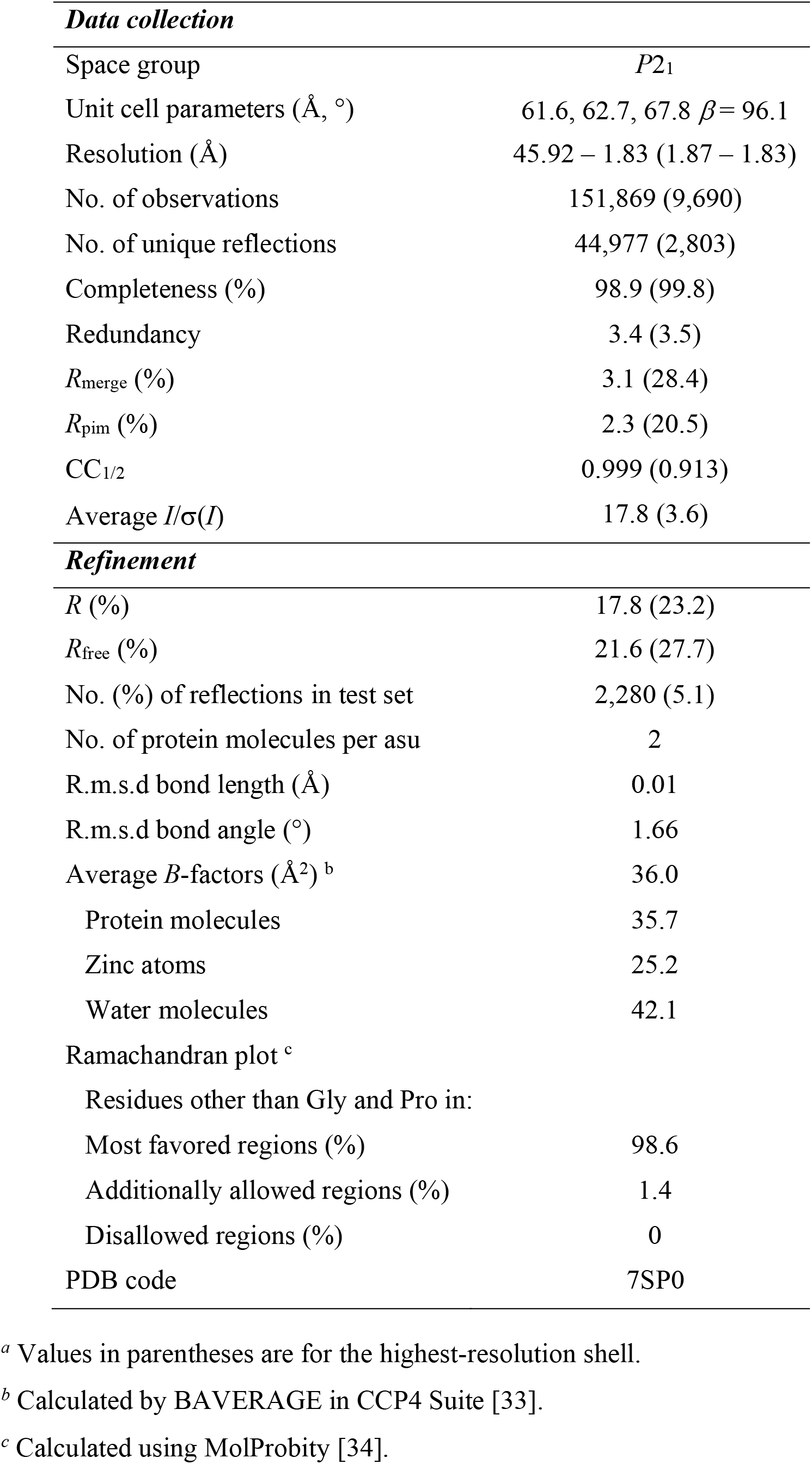
Diffraction data and refinement statistics ^*a*^.

### Structure solution and refinement

The crystal structure was refined using the Zn-SFPQ complex structure (PDB code 6OWJ [19]) as the initial model after removing all non-protein atoms and mutating L534 to alanine. Iterative model building with COOT [31] and refinement with REFMAC5 [32] within the CCP4 suite [33] was carried out. The final model consisted of two chains of SFPQ (residues 283 – 535 in Chain A and residues 290 – 368, 371 – 535 in Chain B), two Zn atoms (one of the two with half occupancy), and 277 water molecules. The quality of the model was validated using MOLPROBITY[34]. The refinement statistics are included in **Table 1**. The atomic coordinates have been deposited in the Protein Data Bank as entry 7SP0.

### Zinc treatment and SFPQ localization assay

Primary cortical neurons were prepared from embryonic day 18 rat pups as described previously [35]. Neurons were transfected at days *in vitro* (DIV) 13 using Lipofectamine 2000 (Invitrogen). The next day, they were treated with either 100 μM ZnCl_2_ for 4 h or water (vehicle control), and subsequently fixed with 4% paraformaldehyde/4% sucrose solution in PBS. Following extensive washes, the neurons were stained with anti-MAP2 antibody (M3696, Sigma-Aldrich) before mounting them onto glass slides using ProLong Diamond anti-fade mounting medium with DAPI (Invitrogen). Slides were imaged on a Zeiss Axio Imager epifluorescence microscope. The fractions of neurons containing nuclear or cytoplasmic GFP-SFPQ aggregates over total transfected neurons were quantified and normalized to the GFP-SFPQ WT without ZnCl_2_ treatment group.

### Dendritic spine analysis

Neurons were co-transfected with the structural marker td-Tomato and pEGFP-SFPQ, either WT, N533H or L534I mutants for 48 h. Fixed neurons were imaged with a 100X oil-immersion objective on an inverted Diskovery spinning disk confocal microscope. Secondary dendrites of co-transfected neurons were randomly imaged as z-stacks with a 0.4 μm step size over a range of approximately 5-10 μm. Spine analysis was performed on raw image stacks (z-plane) using ImageJ software. Spine density was calculated as the total number of spines per sum of all measured dendritic lengths for each neuron. All clear protrusions from the dendrite, irrespective of their orientation relative to the imaging plane, were included in the analyses.

### Surface staining assay

To determine the level of AMPARs on the plasma membrane of primary neurons, we performed an antibody-feeding assay as previously described [23]. Neurons were co-transfected with pRK5-myc-GluA1 or pRK5-myc-GluA2 with pEGFP alone or pEGFP-SFPQ, either WT, N533H or L534I mutants, for 48 h. Surface AMPARs were labeled by incubating live neurons with mouse anti-myc antibody (MCA2200, BioRad) for 30 min at 4°C prior to 10 min fixation in ice-cold parafix solution (4% paraformaldehyde, 4% sucrose in PBS). Following cell permeabilization (0.25% Triton X-100 in PBS, 10 min) and blocking (10% normal goat serum, 1 h) at room temperature, total myc-GluA1/2 was labeled with rabbit anti-myc antibody (71D10, Cell Signaling Technology) at 4°C overnight. The surface and total myc-GluA1 or myc-GluA2 were subsequently visualized by Alexa-568-conjugated anti-mouse and Alexa-647-conjugated anti-rabbit secondary antibodies, respectively. Images were collected with a 63X oil-immersion objective on a Zeiss LSM510 confocal microscope. Fluorescence intensities were quantified using ImageJ software (NIH) for surface and total receptors. Data were expressed as the surface/total AMPAR ratio.

## RESULTS

### SFPQ N533H and L534I display higher zinc-binding affinities than WT protein

We have previously identified an intermolecular zinc-coordinating center within the SFPQ DBHS domain that comprises His-483, His-528, His-530, and the carbonyl group of Leu-535 (**Fig. 1A, B**). The identification of two fALS-associated *SFPQ* missense variants in the proximity of the SFPQ zinc-coordinating center (N533H and L534I) suggests that they may affect the affinity of SFPQ to zinc. In particular, the additional histidine residue in the N533H mutant may be involved in zinc coordination, thereby enhancing the affinity of SFPQ to zinc. To test this hypothesis, we performed a competitive zinc-binding assay on the purified SFPQ DBHS domain containing the N533H or L534I mutations using the fluorescent zinc indicator, Fluozin-3 [19]. The calculated affinity (IC_50_) of SFPQ N533H (0.25 μM) and L534I (0.54 μM) binding to zinc was approximately 5.8- and 2.7-fold higher than that of the WT protein (1.45 μM), respectively (**Fig. 1C** and **Suppl. Table S2**), confirming that these fALS-associated mutations cause significant increases in SFPQ zinc-binding affinity. Although the effect of SFPQ N533H was expected, it was surprising to observe the increased zinc-binding affinity for the L534I mutant given the subtle nature of the substitution. It is also noteworthy that the slope of fluorescence reduction was significantly steeper for SFPQ L534I than SFPQ WT, whereas that of SFPQ N533H was modest, represented by Hill slopes of -25.1 (L534I), -12.4 (N533H), and -5.9 (WT) (**Fig. 1C** and **Suppl. Table S2**). The crystal structure of SFPQ L534I in complex with zinc described below partly explains this cooperativity.

### The crystal structure of SFPQ L534I reveals a secondary zinc-binding site

To gain insights into the structural basis of the enhanced zinc-binding affinity of the fALS-associated SFPQ mutant, we solved the crystal structure of the SFPQ dimerization domain (residue 276-535) containing the L534I mutation in complex with Zn(II) (**Fig. 2A**). The approach to produce crystals of SFPQ WT in complex with Zn(II) was the same as that used previously [19]. Briefly, the DNA-binding domain of retinoic X receptor α (RXRα), which contains two zinc finger domains, was incubated with the SFPQ L534I mutant protein (1:1 molar ratio) and crystallized by the hanging-drop vapor diffusion method at 20°C [19]. The overall structure of SFPQ L534I in complex with Zn(II) is similar to that of SFPQ WT with the root mean square deviation (r.m.s.d.) of 0.28 Å between them (496 common Cα superposed) (**Fig. 2A** and **Suppl. Fig. S1**). Consistent with SFPQ WT, the intermolecular interaction mediated by zinc also results in infinite polymerization of SFPQ L534I (**Fig. 2B, C**). Given the subtle nature of the substitution from leucine to isoleucine, these observations were expected. However, the structural refinement of SFPQ L534I revealed a surprising additional metal density, which also mediates the intermolecular interaction of SFPQ (**Fig. 2B, C**). The identity of this metal ion was confirmed as zinc by calculating the anomalous difference Fourier maps from diffraction data collected at energies near the zinc X-ray absorption edge: 9760 eV (*f”*_Zn_ = 3.77 e^-^) and 9560 eV (*f”*_Zn_ = 0.57 e^-^) (**Fig. 2D, E** and **Suppl. Fig. S2**).

**Fig. 2.**
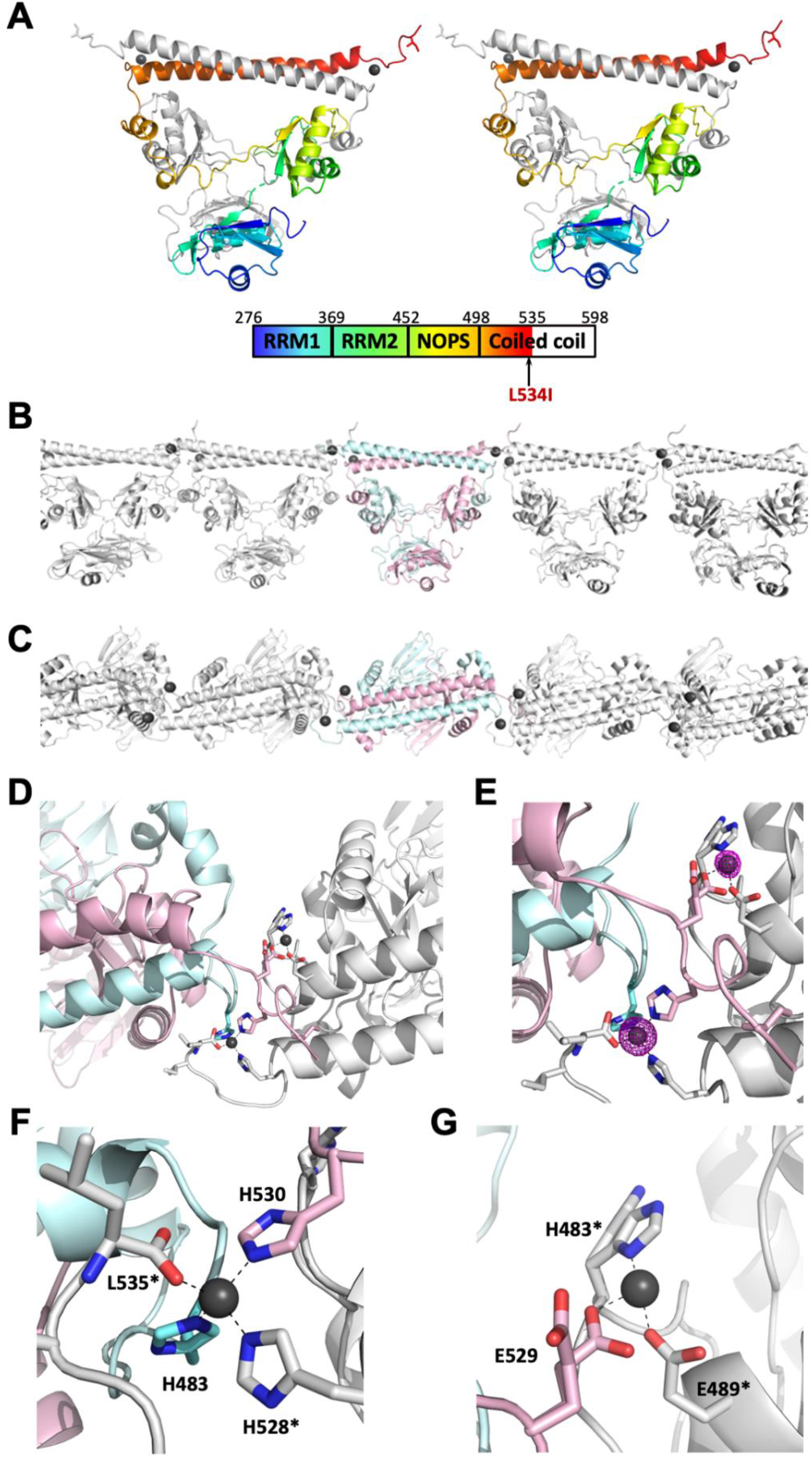
The crystal structure of the SFPQ L534I mutant reveals a second zinc-binding site. **A** Stereo view of the dimerization domain of the human SFPQ (residues 276 – 535) L534I mutant in complex with zinc in cartoon presentation with Chain A in gray and Chain B in the same color scheme as in the schematic domain organization below. The Zn atoms are shown as black spheres. **B, C** Zinc-mediated infinite polymerization observed in the crystal structure of SFPQ L534I from the side (**B**) and top (**C**) views. Chains A and B are shown in cyan and pink, respectively, while the neighboring symmetry-related dimers are shown in gray. **D** Zinc centers in the crystal structure of the SFPQ L534I mutant. The intermolecular interaction mediated by two zinc atoms is shown with Chain A and Chain B in cyan and pink, respectively, while the neighboring dimer (symmetry operator: x, y, z – 1) is shown in gray. **E** Close-up view of the zinc centers with overlaid anomalous difference Fourier map (|*F*^+^| – |*F*^-^|, contoured at 7 σ) from the diffraction data collected at 9760 eV near the Zn X-ray absorption edge (*f”*_Zn_ = 3.77 e^-^) for zinc identification. The positions of the L534I mutation are shown with the side chain of Ile in stick presentation. **F** Zinc center 1 is coordinated by H483 (Chain A), H530 (Chain B), H528* (Chain A of neighboring dimer), and carbonyl oxygen of L535* (Chain A of neighboring dimer). The asterisk (*) denotes a symmetry-related dimer (x, y, z – 1). **G** The partially occupied zinc center 2 (occupancy of 0.5) is coordinated by E529 (Chain B), H483* and E489* (Chain B of neighboring dimer).

A closer inspection of the two zinc centers showed that the first zinc coordinating center (Zn center 1) is almost indistinguishable from that of SFPQ WT, coordinated by His A483, His B530, His A*528 and a carbonyl oxygen of Leu A*535, where the asterisk (*) denotes a symmetry-related dimer (x, y, z – 1) (**Fig. 2F**). The Zn^2+^ ion in this center is fully occupied with a comparable *B*-factor value (22.4 Å^2^) to those of ligating atoms (average *B*-factor of 20.9 Å^2^). The anomalous signal from the second center (Zn center 2) is, however, significantly lower than that of Zn center 1 (**Fig. 2E**), indicating that this site is not fully occupied. The second zinc coordination center is composed of three amino acid residues: Glu B*489, Glu B529, and His B*483 (symmetry operator: x, y, z – 1). Further supporting the partially occupied zinc, the side chain of two out of three of these amino acids is also observed in two different conformations (**Fig. 2G**). The side chains of Glu B529 and His B*483 are in the ‘in’ position coordinating zinc, whereas in the absence of zinc, they take on the ‘out’ conformation. Zinc in the second center is modeled with the occupancy of 0.5 in the final structure with a comparable *B*-factor value (28.1 Å^2^) to those ligating atoms (average *B*-factor of 26.8 Å^2^). The fourth zinc-ligating atom is likely to be a partially occupied solvent atom; however, it could not be modeled in the final structure due to its low electron density (**Suppl. Fig. S2**). Taken together, the crystal structure of SFPQ L534I in complex with zinc reveals an additional zinc-binding site, providing the structural basis for the apparent increase in zinc-binding affinity in this mutant.

### fALS-associated N533H and L534I mutants promote SFPQ cytoplasmic aggregation in neurons

Mislocalization and cytoplasmic aggregation of nuclear SFPQ are associated with the pathogenesis of ALS [15, 16]. We have previously demonstrated that the application of ZnCl_2_ induces the cytoplasmic accumulation and aggregation of SFPQ in cultured neurons [19]. To determine the effect of N533H and L534I mutations on SFPQ localization, we transiently transfected primary cortical neurons with DNA constructs that encode GFP-SFPQ, either WT, N533H or L534I for 24 h. Under basal conditions, overexpression of GFP-SFPQ N533H and L534I significantly enhanced the propensity to form cytoplasmic aggregates in primary cortical neurons compared to the WT protein (**Fig. 3A–C**). As expected, the addition of 100 μM ZnCl_2_ to the culture medium for 4 h significantly increased the number of neurons with cytoplasmic GFP-SFPQ aggregates in the somatodendritic regions (**Fig. 3D**). However, ZnCl_2_ treatment did not result in a further increase in the proportion of neurons containing GFP-SFPQ cytoplasmic aggregates when transfected with the N533H and L534I mutants, suggestive of an occlusion effect (**Fig. 3D**). Together, these data indicate that the increase in zinc-binding affinity accounts for the elevation in the number of cytoplasmic aggregates observed in neurons that express these two fALS-associated SFPQ mutants.

**Fig. 3.**
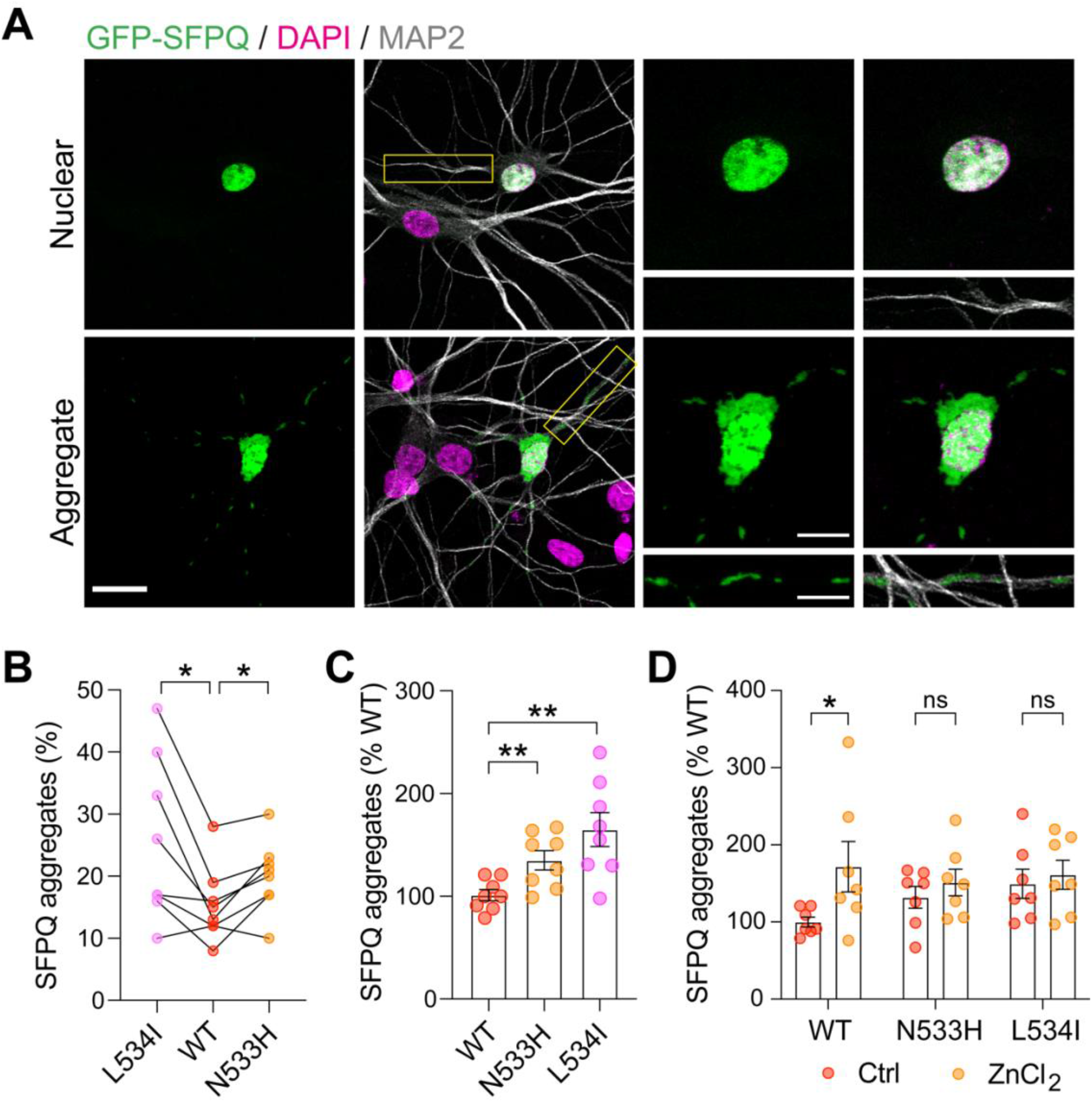
fALS-associated N533H and L534I mutants promote the cytoplasmic aggregation of SFPQ. **A** Representative confocal images of primary cortical neurons expressing GFP-SFPQ (green) exhibiting nuclear localization (top panel) or cytoplasmic aggregates in the somatodendritic regions (bottom). Higher-magnification images are shown on the right panels. Scale bars, 50 μm or 10 μm (inset). **B** Quantification of the fraction of neurons with GFP-SFPQ cytoplasmic aggregates within individual experiments (Wilcoxon matched-pairs t-test, \**p*<0.05, N=5 independent experiments). **C** Quantification of the fraction of neurons with GFP-SFPQ cytoplasmic aggregates normalized to the value of the WT group. Data represent mean ± SEM (Welch’s unpaired t-test, \*\**p*<0.01, N=5 independent experiments). **D** Zinc-induced SFPQ aggregation is occluded in neurons expressing fALS-associated mutants. Primary cortical neurons expressing GFP-SFPQ, either WT, N533H or L534I mutants, were treated with 100 μM ZnCl_2_ for 4 h. Data represent mean ± SEM, normalized to the WT group (two-way ANOVA, Sidak’s post-hoc multiple comparison test, \**p*<0.05, ns=not significant, N=3 independent experiments). Each data point was derived from one coverslip, which contained an average of 113 transfected neurons.

### Cytoplasmic SFPQ aggregation causes a reduction in the level of surface GluA1 expression

Our observation that SFPQ cytoplasmic aggregates are largely localized in the soma and dendrites raises the question of whether these protein aggregates affect dendritic functions in primary neurons. To investigate this, we first examined the effect of SFPQ overexpression on the density of dendritic spines of neurons that co-expressed a structural marker td-Tomato and GFP or GFP-SFPQ, either WT, N533H or L534I (**Fig. 4A**). Interestingly, overexpression of GFP-SFPQ caused a significant reduction in the number of dendritic spines compared with those expressing soluble GFP alone (**Fig. 4A, B**). This effect was observed in all neurons containing nuclear or aggregated GFP-SFPQ (**Fig. 4A, B**). However, there was no significant difference between GFP-SFPQ WT or fALS mutants (**Fig. 4C**), suggesting that these mutations play no role in regulating the density of dendritic spines in primary neurons.

**Fig. 4.**
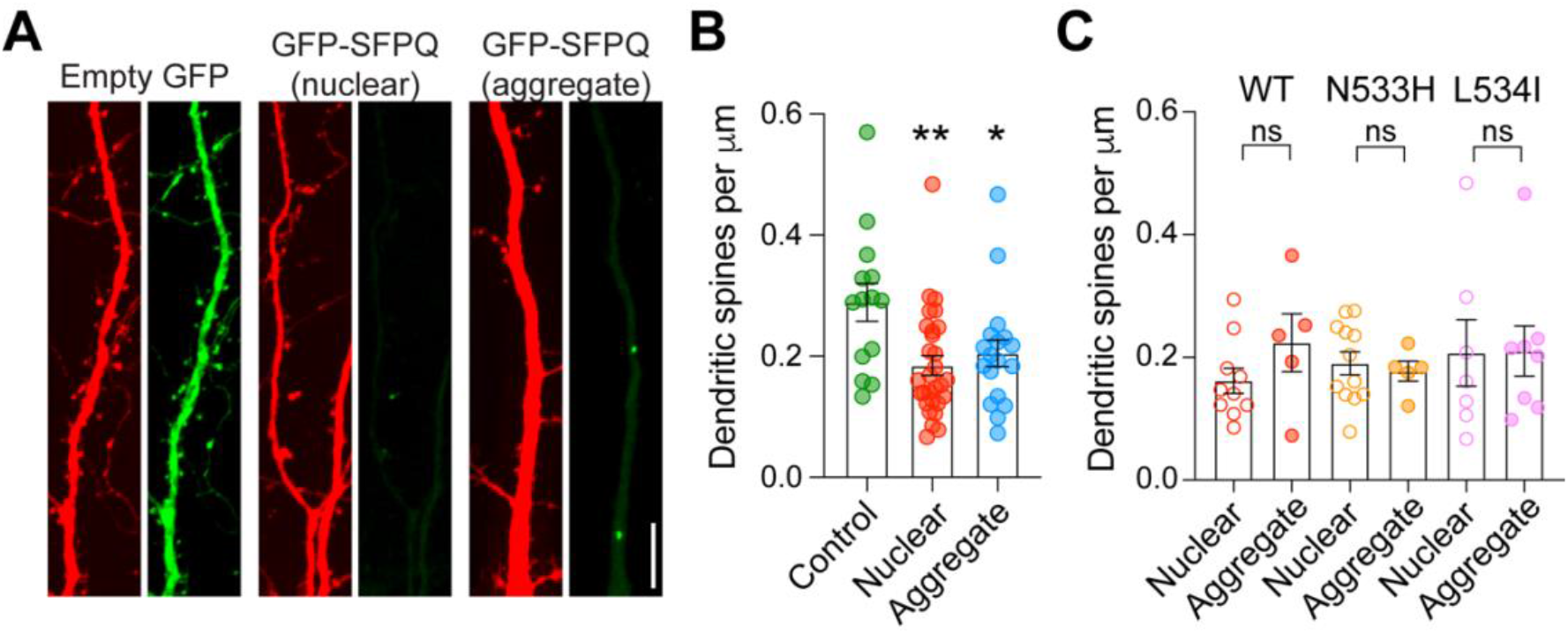
Overexpression of GFP-SFPQ reduces dendritic spine density, a phenotype that is not affected by fALS-associated *SFPQ* variants. **A** Confocal images of dendritic segments of neurons co-expressing the structural marker td-Tomato (red) and soluble GFP (left panels) or GFP-SFPQ (green) that exhibit nuclear localization (middle panels) or cytoplasmic aggregates in the somatodendritic regions (right panels). Scale bar, 10 μm (inset). **B** Quantification of dendritic spine density from neurons that express soluble GFP, nuclear GFP-SFPQ or GFP-SFPQ aggregates. Data are presented as mean ± SEM (one-way ANOVA, Tukey’s post-hoc multiple comparison test, \**p*<0.05 \*\**p*<0.01; GFP, n=14 neurons; nuclear, n=29 neurons; aggregate, n=18 neurons, N=2 independent experiments). **C** Quantification of dendritic spine density from neurons expressing GFP-SFPQ WT, N533H or L534I mutants. Data are presented as mean ± SEM (Welch’s unpaired t-test, ns=not significant; N=2 independent experiments).

Given that excitotoxicity and altered cortical excitability are pathophysiological features associated with ALS [36], we next investigated the effect of the N533H and L534I mutants on the surface expression of the GluA1 and GluA2 subunits of AMPARs, which mediate the majority of fast excitatory synaptic transmission in the mammalian central nervous system [37]. Primary neurons were transfected with myc-GluA1 or myc-GluA2 reporter constructs with plasmids that encode GFP or GFP-SFPQ, either WT, N533H or L534I, followed by a surface staining assay with anti-myc antibodies. Neurons that expressed nuclear GFP-SFPQ, regardless of its genotype, had normal levels of GluA1-containing AMPARs on the plasma membrane compared to those that expressed soluble GFP alone (**Fig. 5A, B**). However, cytoplasmic SFPQ aggregates caused a significant decrease in the expression of GluA1-containing AMPARs on the plasma membrane of neurons that overexpressed GFP-SFPQ WT, N533H or L534I **(Fig. 5A–C)**. This effect was specific to GluA1 as cytoplasmic SFPQ aggregates did not affect the expression of surface GluA2-containing AMPARs in primary neurons (**Fig. 5D, E**). Collectively, our results indicate that the fALS-associated *SFPQ* variants, which promote the propensity of SFPQ to form cytoplasmic aggregates, may alter neuronal excitability by changing the subunit composition of neuronal AMPARs.

**Fig. 5.**
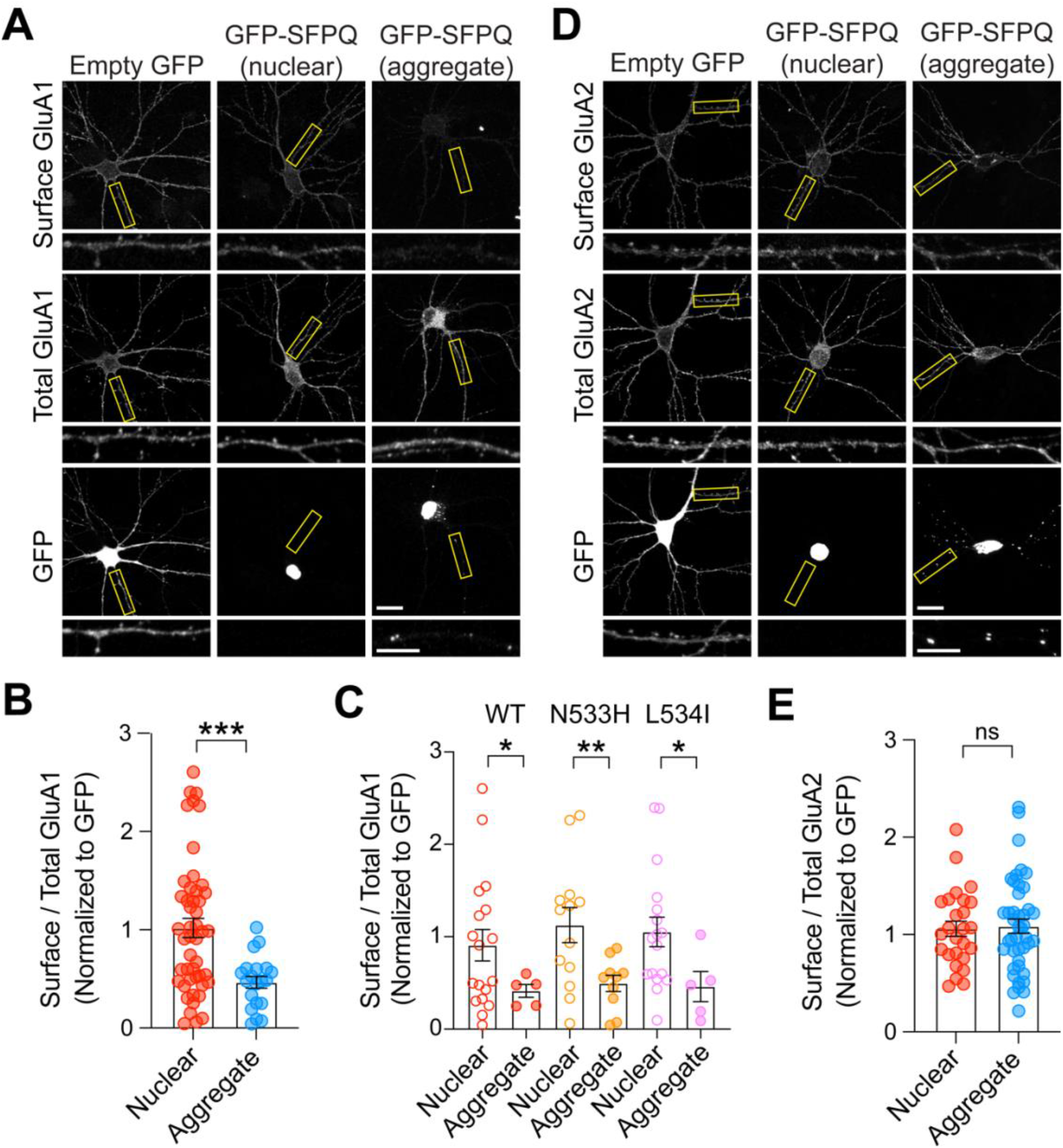
Cytoplasmic SFPQ aggregates selectively reduce the levels of GluA1- but not the GluA2-containing AMPARs on the plasma membrane. **A** Primary cortical neurons were co-transfected with plasmids encoding myc-GluA1 and GFP or GFP-SFPQ. Representative images of surface and total myc-GluA1 in neurons expressing soluble GFP, nuclear GFP-SFPQ or GFP-SFPQ cytoplasmic aggregates. Zoomed images are dendrites from the boxed regions. Scale bars, 20 μm or 10 μm (enlarged images). **B** Quantification of the surface/total GluA1 ratio normalized to the value of GFP-expressing neurons. Data are presented as mean ± SEM (Welch’s unpaired t-test, \*\*\**p*<0.001; nuclear, n=48 neurons; aggregate, n=20 neurons, from two independent experiments). **C** Quantification of the surface/total GluA1 ratio from neurons expressing GFP-SFPQ WT, N533H or L534I mutants, normalized to the value of GFP-expressing neurons. Data are presented as mean ± SEM (Welch’s unpaired t-test, \**p*<0.05 \*\**p*<0.01; N=2 independent experiments). **D** Confocal images of primary neurons co-expressing myc-GluA2 and soluble GFP, nuclear GFP-SFPQ or GFP-SFPQ cytoplasmic aggregates. Scale bars, 20 μm or 10 μm (enlarged images). **E** Cytoplasmic SFPQ aggregation does not affect the expression of surface GluA2-containing AMPARs. Data represent mean ± SEM (Welch’s unpaired t-test, ns=not significant; nuclear, n=26 neurons; aggregate, n=43 neurons, from two independent experiments).

## DISCUSSION

SFPQ is a ubiquitous nuclear RBP that is highly expressed in the brain, with multiple roles in transcriptional regulation, alternative splicing, mRNA transport, paraspeckle formation and RNA metabolism [1, 2]. Nuclear depletion and cytoplasmic accumulation of SFPQ have been linked to the pathophysiology of neurodegenerative diseases, including Alzheimer’s disease [38] and ALS [15, 16]. Despite this, our understanding of how SFPQ mislocalizes into the cytoplasm and forms protein aggregates is very limited. Our previous work has demonstrated the role of zinc in inducing SFPQ polymerization *in vitro* and the formation of SFPQ cytoplasmic aggregates in primary neurons [19]. In the present study, we characterize two fALS-associated *SFPQ* variants that result in amino acid substitutions (N533H and L534I) in the proximity of the SFPQ zinc-coordinating center. We found that both mutants enhance zinc-binding affinity and the propensity to form cytoplasmic aggregates in the somatodendritic region of primary neurons.

### Structural basis of enhanced zinc-binding affinity in SFPQ L534I

The identification of a secondary zinc-binding site in the SFPQ L534I structure explains the apparent increase in zinc-binding affinity. It also explains the higher Hill slope measured in this variant in comparison to the WT protein. Given that there is more than one zinc-binding site in SFPQ L534I, we fitted the data from the zinc-binding assays with the IC_50_ model instead of the one site non-linear fit in our previous study with SFPQ WT [19]. This raises the question as to which structural changes accommodate the second zinc-binding site in this variant. An overall structural comparison between the L534I mutant with SFPQ WT in complex with zinc reveals no obvious differences that can explain the increased zinc-binding of SFPQ L534I (**Suppl. Fig. S1**). The main structural deviations are concentrated at the end of N-terminal region upstream of the structured RRM1 due to lack of crystal contacts. Part of the NOPS domain in Chain A (residues 449 – 477) shows higher r.m.s.d. values than the mean value of 0.28 Å with greatest Cα r.m.s.d. of 1.0 Å (Met-469). However, how these subtle displacements increase the Zn^2+^-binding affinity of SFPQ L534I is not clear.

This led us to revisit the crystal structure of SFPQ WT in complex with zinc. The anomalous difference Fourier maps calculated from the diffraction data collected at the high Zn energy wavelength at 9760 eV clearly show the difference between WT and the L534I mutant (**Suppl. Fig. S3**). In the position corresponding to Zn center 2, there is a small anomalous signal that accounts for a possible weakly bound Zn(II) in SFPQ WT. However, it is significantly weaker than that of SFPQ L534I, reinforcing our observation that this mutation enhances the zinc-binding affinity of SFPQ in solution. The final model of SFPQ WT has a water molecule in this site with a *B*-factor value of 29.5 Å^2^, while Oε1 of Glu-489, a potential hydrogen donor to this water molecule, has a comparable *B*-factor value of 23.3 Å^2^. Although it is subtle, the substitution of leucine to isoleucine may affect the dynamics of SFPQ in solution, thereby facilitating the increased zinc-binding affinity. However, the precise underlying molecular mechanism requires further investigation.

At present, we are not able to ascertain if the additional histidine in SFPQ N533H is indeed replacing the carbonyl oxygen atom of Leu-535 in the zinc-coordinating center in the absence of the zinc-bound SFPQ N533H crystal structure. However, the fact that SFPQ N533H (0.25 μM) displays a higher zinc-binding affinity than SFPQ L534I (0.54 μM), and the failure to crystallize this mutant under the same conditions as SFPQ WT and L534I may indicate that SFPQ N533H has a different zinc center.

### Cytoplasmic SFPQ aggregates selectively reduce the expression of GluA1-containing AMPARs

Excitotoxicity is a major cause of neuronal death in a number of neurodegenerative conditions, including ALS [39], with altered cortical excitability and synaptic dysfunction being identified as early pathophysiological features in this disease [36, 40]. Emerging evidence has demonstrated attenuated synaptic function in neurons expressing ALS-associated mutant proteins, particularly alterations in their dendritic arbor complexity, dendritic spine density and the level of postsynaptic ionotropic AMPARs [41-43]. We found that overexpression of GFP-SFPQ significantly downregulates the number of dendritic spines in primary cortical neurons compared to those that express soluble GFP. Interestingly, this effect was observed in all neurons with nuclear or cytoplasmic SFPQ aggregates. One plausible mechanism may involve the sequestration of an important SFPQ interacting partner FUS, leading to its loss of function and consequently, a deficit in dendritic spine morphogenesis [44-46]. Alternatively, overexpression of SFPQ can dysregulate the transcription of long genes, many of which are involved in neuronal development and neurite outgrowth [5]. Although SFPQ overexpression-induced loss of dendritic spines is not affected by fALS-associated *SFPQ* variants, our data suggest that SFPQ is an important regulator of dendritic spine formation and/or maturation in excitatory neurons, a finding which warrants further investigation.

AMPARs are assembled as two identical heterodimers of GluA1-4 subunits that form functional glutamate-gated ion channels. The presence of the GluA2 subunit renders AMPARs impermeable to Ca^2+^. Alterations in AMPAR subunit composition have been reported in human post-mortem tissues of ALS patients [47, 48] and in various ALS models that overexpress ALS-associated proteins, including TDP-43, FUS and C9ORF72 [44, 49-51]. Consistent with this notion, we found that neurons that contain cytoplasmic SFPQ aggregates have significantly reduced surface expression of GluA1-containing AMPARs compared to those that express nuclear SFPQ. This effect is specific to the GluA1 subunit as the aggregation of SFPQ does not affect the GluA2 subunit. Interestingly, the two fALS-associated *SFPQ* variants do not directly contribute to altering AMPAR subunit composition in neurons per se given that their effects are indistinguishable from those that express SFPQ WT.

ALS-associated alterations in AMPAR subunit composition can occur due to dysregulation of *GRIA* transcripts or inefficient RNA editing of *GRIA2* mRNA that affects AMPAR permeability to Ca^2+^ [39, 52]. However, we posit that cytoplasmic SFPQ aggregates are likely to perturb the trafficking of GluA1-containing AMPARs in primary neurons. Our hypothesis is supported by the fact that SFPQ is present in AMPAR-containing vesicles purified from mouse whole brain lysates [53], and the demonstration that it also interacts with the motor protein KIF5 [11], which has previously been shown to regulate the dendritic transport of such vesicles [54-57]. Importantly, mutations in the KIF5A C-terminal cargo binding domain are associated with fALS [58, 59]. This highlights that disruption in KIF5- mediated transport of not only RNA granules but also neurotransmitter receptors contributes to the pathogenesis of ALS. Although our findings are consistent with the general idea of a perturbation of AMPAR subunit expression as a feature of ALS pathology, cytoplasmic SFPQ aggregates may not cause an increase in the expression of GluA2-lacking Ca^2+^-permeable AMPARs, at least in cortical neurons, that are thought to mediate the excessive Ca^2+^ influx and death of motor neurons [50, 60, 61]. However, it remains to be determined whether a similar effect is observed in motor neurons, given that the heterogeneity of AMPAR subunit expression has been observed in different regions of the brain and might be dependent on genetic causes of the disease [47].

In conclusion, our current study provides the structural basis to explain the high propensity of fALS-associated *SFPQ* variants to form cytoplasmic aggregates through enhanced zinc-binding affinity, leading to a loss in SFPQ function in neurons. Although, it is well established that genetic variants (familial or sporadic) are the main components that predispose (or cause) an individual to develop ALS [62], the molecular mechanisms underlying pathogenesis and disease progression are likely to differ, depending on the mutations. The *SOD1* gene, which encodes the Cu^2+^/Zn^2+^ superoxide dismutase 1 that protects neurons from oxidative stress, is often mutated in fALS patients. Mechanistically, ALS-associated mutations alter the metal binding capacity of SOD1, leading to protein misfolding and aggregation. Accordingly, delivering zinc to mutant SOD1 exerts protective therapeutic effects in the SOD1-G37R ALS mouse model [63]. On the contrary, our findings predict that zinc supplementation would exacerbate SFPQ aggregation and therefore have negative impacts on neuronal function in patients carrying the N533H or L534I mutation. Thus, the outcomes of our study highlight the potential importance of personalized medicine – different mutations may require different interventions – in the treatment of ALS patients with distinct genetic variants.

## Acknowledgements

We thank the Macromolecular Crystallography beamline staff at the Australian Synchrotron (Victoria, Australia) for their professional support. We thank Rowan Tweedale for editing the manuscript. Imaging was performed at the Queensland Brain Institute’s Advanced Microscopy Facility.

## Abbreviations

ALS: amyotrophic lateral sclerosis
AMPA: α-amino-3-hydroxy-5-methyl-4- isoxazoleproprionic acid
AMPAR: AMPA receptor
DBHS: Drosophila behavior human splicing
fALS: familial ALS
FUS: fused in sarcoma
RBP: RNA-binding protein
r.m.s.d.: root mean square deviation
RRM: RNA-recognition motif
RXRα: retinoic X receptor α
SFPQ: splicing factor proline- and glutamine-rich
TDP-43: trans-activation response element DNA-binding protein 43
WT: wild-type.

## REFERENCES

1. Lim YW, James D, Huang J et al (2020) The emerging role of the RNA-binding protein SFPQ in neuronal function and neurodegeneration. Int J Mol Sci 21:7151. https://doi.org/10.3390/ijms21197151

2. Knott GJ, Bond CS, Fox AH (2016) The DBHS proteins SFPQ, NONO and PSPC1: a multipurpose molecular scaffold. Nucleic Acids Res 44:3989–4004. https://doi.org/10.1093/nar/gkw271

3. Lowery LA, Rubin J, Sive H (2007) Whitesnake/sfpq is required for cell survival and neuronal development in the zebrafish. Dev Dyn 236:1347–1357. https://doi.org/10.1002/dvdy.21132

4. Cosker KE, Fenstermacher SJ, Pazyra-Murphy MF et al (2016) The RNA-binding protein SFPQ orchestrates an RNA regulon to promote axon viability. Nat Neurosci 19:690–696. https://doi.org/10.1038/nn.4280

5. Takeuchi A, Iida K, Tsubota T et al (2018) Loss of Sfpq causes long-gene transcriptopathy in the brain. Cell Rep 23:1326–1341. https://doi.org/10.1016/j.celrep.2018.03.141

6. Hewage TW, Caria S, Lee M (2019) A new crystal structure and small-angle X-ray scattering analysis of the homodimer of human SFPQ. Acta Crystallogr F Struct Biol Commun 75:439–449. https://doi.org/10.1107/S2053230X19006599

7. Lee M, Sadowska A, Bekere I et al (2015) The structure of human SFPQ reveals a coiled-coil mediated polymer essential for functional aggregation in gene regulation. Nucleic Acids Res 43:3826–3840. https://doi.org/10.1093/nar/gkv156

8. Huang J, Casas Garcia GP, Perugini MA et al (2018) Crystal structure of a SFPQ/PSPC1 heterodimer provides insights into preferential heterodimerization of human DBHS family proteins. J Biol Chem 293:6593–6602. https://doi.org/10.1074/jbc.RA117.001451

9. Kanai Y, Dohmae N, Hirokawa N (2004) Kinesin transports RNA: isolation and characterization of an RNA-transporting granule. Neuron 43:513–525. https://doi.org/10.1016/j.neuron.2004.07.022

10. Furukawa MT, Sakamoto H, Inoue K (2015) Interaction and colocalization of HERMES/RBPMS with NonO, PSF, and G3BP1 in neuronal cytoplasmic RNP granules in mouse retinal line cells. Genes Cells 20:257–266. https://doi.org/10.1111/gtc.12224

11. Fukuda Y, Pazyra-Murphy MF, Silagi ES et al (2021) Binding and transport of SFPQ-RNA granules by KIF5A/KLC1 motors promotes axon survival. J Cell Biol 220. https://doi.org/10.1083/jcb.202005051

12. Conlon EG, Manley JL (2017) RNA-binding proteins in neurodegeneration: mechanisms in aggregate. Genes Dev 31:1509–1528. https://doi.org/10.1101/gad.304055.117

13. Nussbacher JK, Tabet R, Yeo GW et al (2019) Disruption of RNA metabolism in neurological diseases and emerging therapeutic interventions. Neuron 102:294–320. https://doi.org/10.1016/j.neuron.2019.03.014

14. Loughlin FE, Wilce JA (2019) TDP-43 and FUS-structural insights into RNA recognition and self-association. Curr Opin Struct Biol 59:134–142. https://doi.org/10.1016/j.sbi.2019.07.012

15. Luisier R, Tyzack GE, Hall CE et al (2018) Intron retention and nuclear loss of SFPQ are molecular hallmarks of ALS. Nat Commun 9:2010. https://doi.org/10.1038/s41467-018-04373-8

16. Thomas-Jinu S, Gordon PM, Fielding T et al (2017) Non-nuclear pool of splicing factor SFPQ regulates axonal transcripts required for normal motor development. Neuron 94:322–336 e325. https://doi.org/10.1016/j.neuron.2017.03.026

17. Leal SS, Botelho HM, Gomes CM (2012) Metal ions as modulators of protein conformation and misfolding in neurodegeneration. Coordin Chem Rev 256:2253–2270. https://doi.org/10.1016/j.ccr.2012.04.004

18. Frederickson CJ, Koh JY, Bush AI (2005) The neurobiology of zinc in health and disease. Nat Rev Neurosci 6:449–462. https://doi.org/10.1038/nrn1671

19. Huang J, Ringuet M, Whitten AE et al (2020) Structural basis of the zinc-induced cytoplasmic aggregation of the RNA-binding protein SFPQ. Nucleic Acids Res 48:3356–3365. https://doi.org/10.1093/nar/gkaa076

20. Hozumi I, Yamada M, Uchida Y et al (2008) The expression of metallothioneins is diminished in the spinal cords of patients with sporadic ALS. Amyotroph Lateral Scler 9:294–298. https://doi.org/10.1080/17482960801934312

21. Hozumi I, Hasegawa T, Honda A et al (2011) Patterns of levels of biological metals in CSF differ among neurodegenerative diseases. J Neurol Sci 303:95–99. https://doi.org/10.1016/j.jns.2011.01.003

22. Kanias GD, Kapaki E (1997) Trace elements, age, and sex in amyotrophic lateral sclerosis disease. Biol Trace Elem Res 56:187–201. https://doi.org/10.1007/BF02785392

23. Anggono V, Koc-Schmitz Y, Widagdo J et al (2013) PICK1 interacts with PACSIN to regulate AMPA receptor internalization and cerebellar long-term depression. Proc Natl Acad Sci U S A 110:13976–13981. https://doi.org/10.1073/pnas.1312467110

24. Widagdo J, Fang H, Jang SE et al (2016) PACSIN1 regulates the dynamics of AMPA receptor trafficking. Sci Rep 6:31070. https://doi.org/10.1038/srep31070

25. Zhao Q, Khorasanizadeh S, Miyoshi Y et al (1998) Structural elements of an orphan nuclear receptor-DNA complex. Mol Cell 1:849–861

26. Lim YW, Lee M (2020) Rapid purification method for human SFPQ by implementing zinc-induced polymerization. Protein Expr Purif 171:105626. https://doi.org/10.1016/j.pep.2020.105626

27. Monk IR, Shaikh N, Begg SL et al (2019) Zinc-binding to the cytoplasmic PAS domain regulates the essential WalK histidine kinase of Staphylococcus aureus. Nat Commun 10:3067. https://doi.org/10.1038/s41467-019-10932-4

28. Aragao D, Aishima J, Cherukuvada H et al (2018) MX2: a high-flux undulator microfocus beamline serving both the chemical and macromolecular crystallography communities at the Australian Synchrotron. J Synchrotron Radiat 25:885–891. https://doi.org/10.1107/S1600577518003120

29. Kabsch W (2010) Xds. Acta Crystallogr D66:125–132. https://doi.org/10.1107/S0907444909047337

30. Evans PR, Murshudov GN (2013) How good are my data and what is the resolution? Acta Crystallogr D69:1204–1214. https://doi.org/10.1107/S0907444913000061

31. Emsley P, Lohkamp B, Scott WG et al (2010) Features and development of Coot. Acta Crystallogr D Biol Crystallogr 66:486–501. https://doi.org/10.1107/S0907444910007493

32. Kovalevskiy O, Nicholls RA, Long F et al (2018) Overview of refinement procedures within REFMAC5: utilizing data from different sources. Acta Crystallogr D 74:215–227. https://doi.org/10.1107/S2059798318000979

33. Winn MD, Ballard CC, Cowtan KD et al (2011) Overview of the CCP4 suite and current developments. Acta Crystallogr D67:235–242. https://doi.org/10.1107/S0907444910045749

34. Chen VB, Arendall WB, 3rd, Headd JJ et al (2010) MolProbity: all-atom structure validation for macromolecular crystallography. Acta Crystallogr D66:12–21. https://doi.org/10.1107/S0907444909042073

35. Widagdo J, Chai YJ, Ridder MC et al (2015) Activity-dependent ubiquitination of GluA1 and GluA2 regulates AMPA receptor intracellular sorting and degradation. Cell Rep 10:783–795. https://doi.org/10.1016/j.celrep.2015.01.015

36. Fogarty MJ (2018) Driven to decay: Excitability and synaptic abnormalities in amyotrophic lateral sclerosis. Brain Res Bull 140:318–333. https://doi.org/10.1016/j.brainresbull.2018.05.023

37. Anggono V, Huganir RL (2012) Regulation of AMPA receptor trafficking and synaptic plasticity. Curr Opin Neurobiol 22:461–469. https://doi.org/10.1016/j.conb.2011.12.006

38. Ke Y, Dramiga J, Schutz U et al (2012) Tau-mediated nuclear depletion and cytoplasmic accumulation of SFPQ in Alzheimer’s and Pick’s disease. PLoS One 7:e35678. https://doi.org/10.1371/journal.pone.0035678

39. Van Den Bosch L, Van Damme P, Bogaert E et al (2006) The role of excitotoxicity in the pathogenesis of amyotrophic lateral sclerosis. Biochim Biophys Acta 1762:1068–1082. https://doi.org/10.1016/j.bbadis.2006.05.002

40. Geevasinga N, Menon P, Ozdinler PH et al (2016) Pathophysiological and diagnostic implications of cortical dysfunction in ALS. Nat Rev Neurol 12:651–661. https://doi.org/10.1038/nrneurol.2016.140

41. Handley EE, Pitman KA, Dawkins E et al (2017) Synapse dysfunction of layer V pyramidal neurons precedes reurodegeneration in a mouse model of TDP-43 proteinopathies. Cereb Cortex 27:3630–3647. https://doi.org/10.1093/cercor/bhw185

42. Jiang T, Handley E, Brizuela M et al (2019) Amyotrophic lateral sclerosis mutant TDP-43 may cause synaptic dysfunction through altered dendritic spine function. Dis Model Mech 12. https://doi.org/10.1242/dmm.038109

43. Fogarty MJ, Mu EW, Noakes PG et al (2016) Marked changes in dendritic structure and spine density precede significant neuronal death in vulnerable cortical pyramidal neuron populations in the SOD1(G93A) mouse model of amyotrophic lateral sclerosis. Acta Neuropathol Commun 4:77. https://doi.org/10.1186/s40478-016-0347-y

44. Udagawa T, Fujioka Y, Tanaka M et al (2015) FUS regulates AMPA receptor function and FTLD/ALS-associated behaviour via GluA1 mRNA stabilization. Nat Commun 6:7098. https://doi.org/10.1038/ncomms8098

45. Ishigaki S, Fujioka Y, Okada Y et al (2017) Altered tau isoform ratio caused by loss of FUS and SFPQ function leads to FTLD-like phenotypes. Cell Rep 18:1118–1131. https://doi.org/10.1016/j.celrep.2017.01.013

46. Ishigaki S, Riku Y, Fujioka Y et al (2020) Aberrant interaction between FUS and SFPQ in neurons in a wide range of FTLD spectrum diseases. Brain 143:2398–2405. https://doi.org/10.1093/brain/awaa196

47. Gregory JM, Livesey MR, McDade K et al (2020) Dysregulation of AMPA receptor subunit expression in sporadic ALS post-mortem brain. J Pathol 250:67–78. https://doi.org/10.1002/path.5351

48. Kawahara Y, Kwak S, Sun H et al (2003) Human spinal motoneurons express low relative abundance of GluR2 mRNA: an implication for excitotoxicity in ALS. J Neurochem 85:680–689. https://doi.org/10.1046/j.1471-4159.2003.01703.x

49. Dyer MS, Reale LA, Lewis KE et al (2021) Mislocalisation of TDP-43 to the cytoplasm causes cortical hyperexcitability and reduced excitatory neurotransmission in the motor cortex. J Neurochem 157:1300–1315. https://doi.org/10.1111/jnc.15214

50. Selvaraj BT, Livesey MR, Zhao C et al (2018) C9ORF72 repeat expansion causes vulnerability of motor neurons to Ca<sup>2+</sup>-permeable AMPA receptor-mediated excitotoxicity. Nat Commun 9:347. https://doi.org/10.1038/s41467-017-02729-0

51. Wright AL, Della Gatta PA, Le S et al (2021) Riluzole does not ameliorate disease caused by cytoplasmic TDP-43 in a mouse model of amyotrophic lateral sclerosis. Eur J Neurosci 54:6237–6255. https://doi.org/10.1111/ejn.15422

52. Kawahara Y, Kwak S (2005) Excitotoxicity and ALS: what is unique about the AMPA receptors expressed on spinal motor neurons? Amyotroph Lateral Scler Other Motor Neuron Disord 6:131–144. https://doi.org/10.1080/14660820510037872

53. Peters JJ, Leitz J, Oses-Prieto JA et al (2021) Molecular characterization of AMPA-receptor-containing vesicles. Front Mol Neurosci 14:754631. https://doi.org/10.3389/fnmol.2021.754631

54. Fan R, Lai KO (2021) Understanding how kinesin motor proteins regulate postsynaptic function in neuron. FEBS J. https://doi.org/10.1111/febs.16285

55. Setou M, Seog DH, Tanaka Y et al (2002) Glutamate-receptor-interacting protein GRIP1 directly steers kinesin to dendrites. Nature 417:83–87. https://doi.org/10.1038/nature743

56. Brachet A, Lario A, Fernandez-Rodrigo A et al (2021) A kinesin 1-protrudin complex mediates AMPA receptor synaptic removal during long-term depression. Cell Rep 36:109499. https://doi.org/10.1016/j.celrep.2021.109499

57. Heisler FF, Lee HK, Gromova KV et al (2014) GRIP1 interlinks N-cadherin and AMPA receptors at vesicles to promote combined cargo transport into dendrites. Proc Natl Acad Sci U S A 111:5030–5035. https://doi.org/10.1073/pnas.1304301111

58. Brenner D, Yilmaz R, Muller K et al (2018) Hot-spot KIF5A mutations cause familial ALS. Brain 141:688–697. https://doi.org/10.1093/brain/awx370

59. Nicolas A, Kenna KP, Renton AE et al (2018) Genome-wide analyses identify KIF5A as a novel ALS gene. Neuron 97:1268–1283 e1266. https://doi.org/10.1016/j.neuron.2018.02.027

60. Tateno M, Sadakata H, Tanaka M et al (2004) Calcium-permeable AMPA receptors promote misfolding of mutant SOD1 protein and development of amyotrophic lateral sclerosis in a transgenic mouse model. Hum Mol Genet 13:2183–2196. https://doi.org/10.1093/hmg/ddh246

61. Kawahara Y, Ito K, Sun H et al (2004) Glutamate receptors: RNA editing and death of motor neurons. Nature 427:801. https://doi.org/10.1038/427801a

62. Mejzini R, Flynn LL, Pitout IL et al (2019) ALS genetics, mechanisms, and therapeutics: Where are we now? Front Neurosci 13:1310. https://doi.org/10.3389/fnins.2019.01310

63. McAllum EJ, Roberts BR, Hickey JL et al (2015) Zn II(atsm) is protective in amyotrophic lateral sclerosis model mice via a copper delivery mechanism. Neurobiol Dis 81:20–24. https://doi.org/10.1016/j.nbd.2015.02.023

