## Supplementary Data for "Familial ALS-associated *SFPQ* variants promote the formation of SFPQ cytoplasmic aggregates that reduce surface AMPA receptor expression in primary neurons"

**Supporting Information**

**Table S1.** Data collection statistics for metal identification*

| | **Zn-high energy** | **Zn-low energy** |
|---|---|---|
| Energy (eV) | 9760 | 9560 |
| Space group | $P2_1$ | $P2_1$ |
| Unit cell parameters (Å, °) | 61.9, 62.8, 68.0, $\beta$ = 95.9 | 61.7, 62.7, 67.9 $\beta$ = 96.1 |
| Resolution (Å) | 45.89 – 2.09 (2.16 – 2.09) | 45.94 – 2.14 (2.20 – 2.14) |
| Completeness (%) | 99.4 (94.6) | 99.6 (96.5) |
| Multiplicity | 6.8 (6.7) | 6.0 (5.9) |
| Anomalous completeness (%) | 98.6 (91.3) | 98.2 (93.7) |
| Anomalous multiplicity | 3.5 (3.5) | 3.0 (3.0) |
| $R_{merge}$ (%) | 5.9 (23.8) | 7.0 (26.5) |
| $R_{pim}$ (%) | 2.7 (10.5) | 3.4 (12.8) |
| $CC_{1/2}$ | 0.998 (0.977) | 0.997 (0.965) |
| Average $I/\sigma(I)$ | 14.8 (4.8) | 11.7 (4.2) |

* A single crystal was used to collect X-ray diffraction datasets at the two different wavelengths.

**Table S2.**

$IC_{50}$ values calculated for the interactions between SFPQ and zinc†

| **SFPQ construct** | **$IC_{50}$ (μM)** | Hill slope‡ | $R^2$ |
|---|---|---|---|
| SFPQ-276–598 WT | 1.45 ± 0.06 | -5.84 | 0.97 |
| SFPQ-276–598 N533H | 0.25 ± 0.01 | -12.36 | 0.99 |
| SFPQ-276–598 L534I | 0.54 ± 0.01 | -25.10 | 0.95 |

† Data are presented with the standard deviation of independent experiments ($n$ = 3).

‡ Hill slope, also called a slope factor, is a quantitative indicator of the steepness of dose-response curves. The data were analyzed using the '$log_{10}$[inhibitor] versus response – variable slope' model in Prism (GraphPad Software). The $IC_{50}$ value for zinc-binding affinity was determined by the following formula: Y = *Bottom* + (*Top* – *Bottom*)/(1 + 10^((Log[$IC_{50}$] – Log[inhibitor])*Hill Slope)) where *Top* and *Bottom* represent plateaus of the fluorescence signal (Y axis) in the curve.

**Fig. S1**

**A**

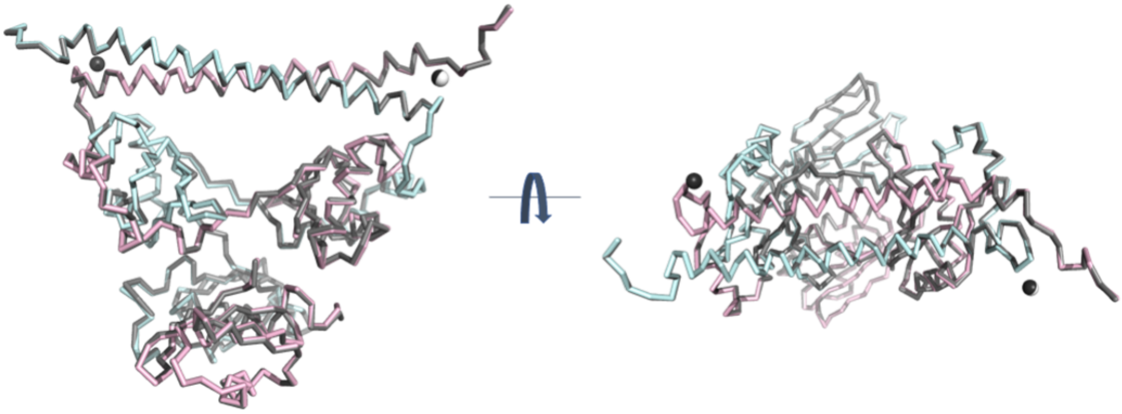

**B**

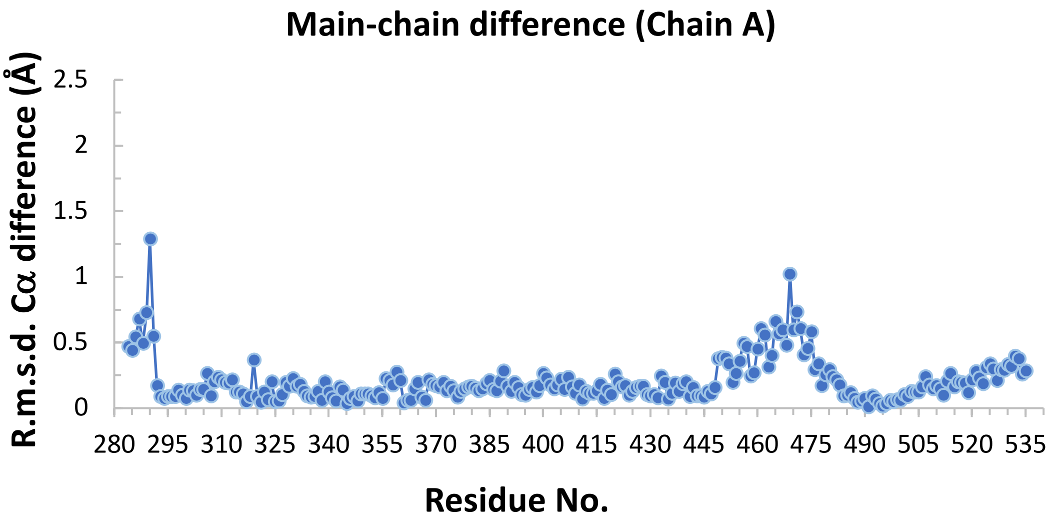
Main-chain difference (Chain A)
R.m.s.d. Cα difference (Å)
2.5
2
1.5
1
0.5
0
280 295 310 325 340 355 370 385 400 415 430 445 460 475 490 505 520 535
Residue No.

**C**

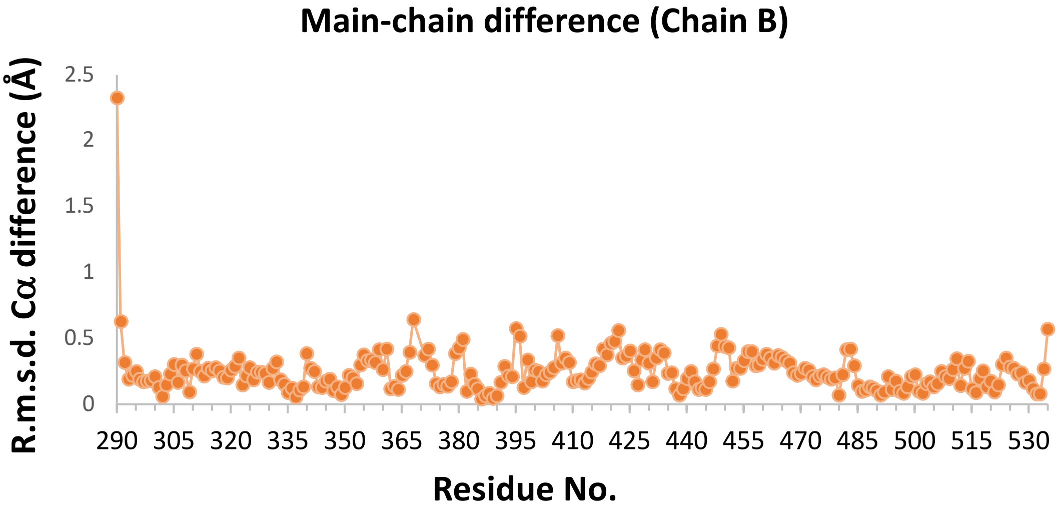
Main-chain difference (Chain B)
R.m.s.d. Cα difference (Å)
2.5
2
1.5
1
0.5
0
290 305 320 335 350 365 380 395 410 425 440 455 470 485 500 515 530
Residue No.

Comparison of the Zn-SFPQ L534I complex (PDB code 7SP0) with the Zn-SFPQ WT (PDB code 6OWJ). **A** Superposition of the Zn-SFPQ L534I complex (67SP0; Chain A in cyan and Chain B in pink with Zn atoms shown as black spheres) and the Zn-SFPQ WT complex (6OWJ in grey with Zn atom shown as white sphere). Both structures are shown in ribbon presentation. **B, C** R.m.s.d. Cα differences of Chain A and Chain B in the two structures.

**Figure S2.**

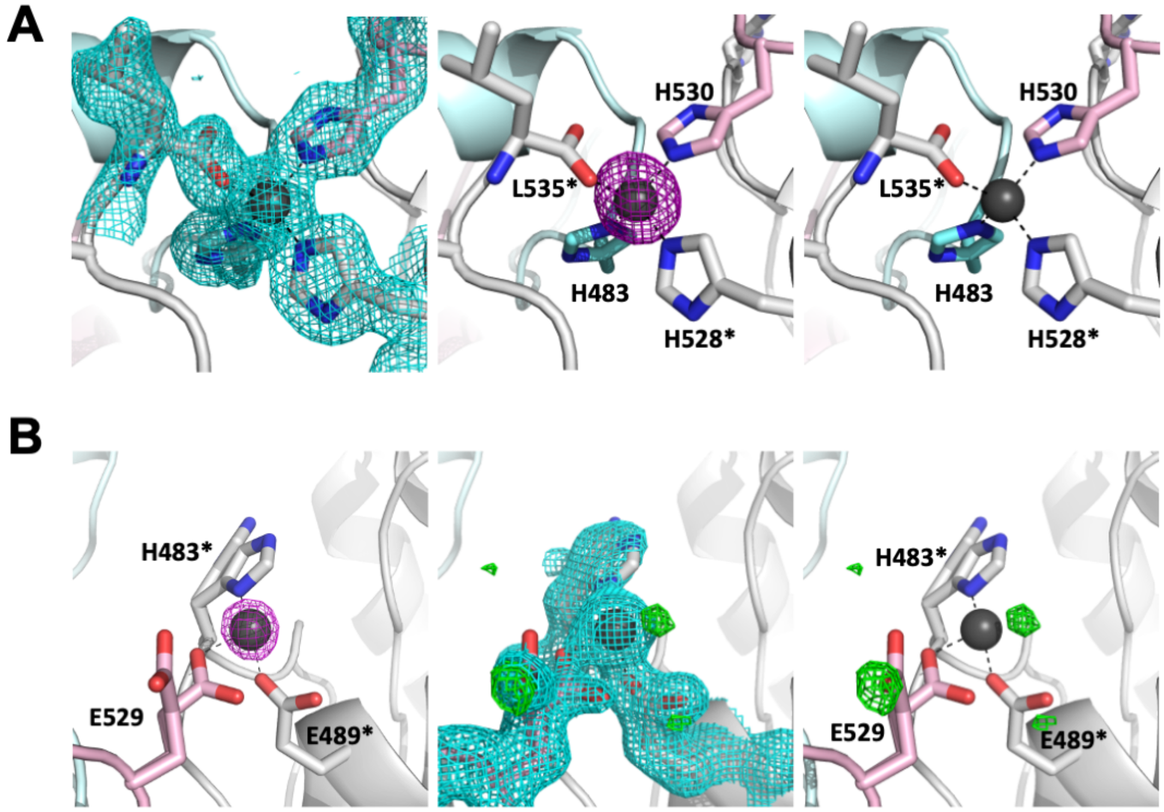


Zn identification via anomalous scattering data collection. **A** Zn center 1 with fully occupied Zn(II). The electron density map ($2F_o - F_c$ map at 1 σ) from the final round of refinement is shown with the final Zn(II)-SFPQ complex model (left panel). Anomalous difference Fourier maps ($|F^+| - |F^-|$, contoured at 7 σ) of the Zn(II)-SFPQ complex from the diffraction data collected at energies near the Zn(II) X-ray absorption edge: (middle panel) 9760 eV ($f''_{Zn}$ = 3.77 $e^-$) and (right panel) 9560 eV ($f''_{Zn}$ = 0.57 $e^-$). **B** Zn center 2 with partially occupied Zn(II) (occupancy of 0.5). Anomalous difference Fourier map ($|F^+| - |F^-|$, contoured at 7 σ) from the diffraction data collected at 9760 eV ($f''_{Zn}$ = 3.77 $e^-$) (left panel). The electron density maps ($2F_o - F_c$ map at 1 σ in cyan; $F_o - F_c$ map at 3.5 σ in green) from the final round of refinement are shown (middle panel). The fourth Zn-ligating atom is likely a partially occupied solvent atom, displayed by the positive $F_o - F_c$ density near Zn (right panel). Carbon atoms in Chain A and Chain B are colored in cyan and pink, respectively, while those from the neighboring dimer (symmetry operator, $x, y, z - 1$) are shown in gray.

**Figure S3.**

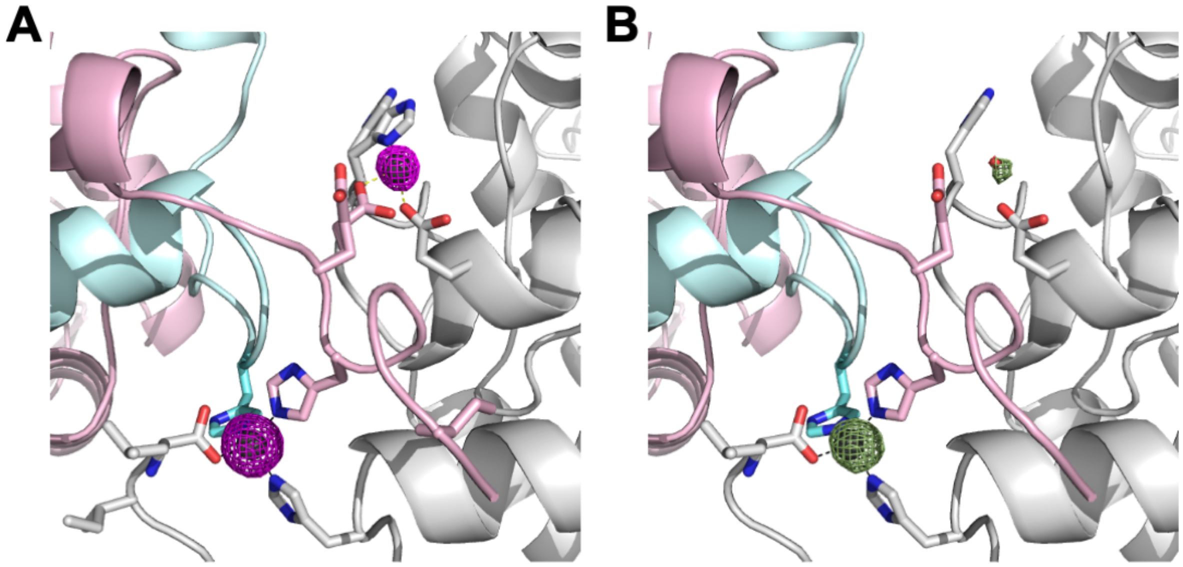


Comparison of the Zn centers of SFPQ L534I with SFPQ WT (PDB code 6OWJ) [19]. Anomalous difference Fourier maps ($|F^+| - |F^-|$, contoured at 7 σ) from the diffraction data collected at 9760 eV ($f''_{Zn}$ = 3.77 e$^-$) are displayed in purple mesh for SFPQ L534I (**A**) and in dark green mesh for SFPQ WT (**B**).
